# Robust metabolic transcriptional components in 34,494 patient-derived samples and cell lines

**DOI:** 10.1101/2020.10.01.321950

**Authors:** V.C. Leeuwenburgh, C.G. Urzúa-Traslaviña, A. Bhattacharya, M.T.C. Walvoort, M. Jalving, S. de Jong, R.S.N. Fehrmann

## Abstract

Patient-derived expression profiles of cancers can provide insight into transcriptional changes that underlie reprogrammed metabolism in cancer. These profiles represent the average expression pattern of all heterogeneous tumor and non-tumor cells present in biopsies of tumor lesions. Therefore, subtle transcriptional footprints of metabolic processes can be concealed by other biological processes and experimental artifacts. We, therefore, performed consensus Independent Component Analyses (c-ICA) with 34,494 bulk expression profiles of patient-derived tumor biopsies, non-cancer tissues, and cell lines. c-ICA enabled us to create a transcriptional metabolic landscape in which many robust metabolic transcriptional components and their activation score in individual samples were defined. Here we demonstrate how this landscape can be used to explore associations between the metabolic transcriptome and drug sensitivities, patient outcomes, and the composition of the immune tumor microenvironment. The metabolic landscape can be explored at http://www.themetaboliclandscapeofcancer.com.

## INTRODUCTION

Reprogrammed energy metabolism is a hallmark of cancer (Hanahan and Weinberg, 2011). Metabolic reprogramming supports the survival, proliferation, and maintenance of cancer cells by ensuring sufficient biosynthetic capacity, redox potential, and energy (Pavlova and Thompson, 2016; Vazquez et al., 2016). Additionally, metabolic reprogramming enables tumor cells to adapt to challenging microenvironmental conditions, such as hypoxia and low nutrient availability, and become resistant to cancer treatment (Huang et al., 2014; Viale and Draetta, 2016). Moreover, metabolic reprogramming of cancer cells influences the composition and function of immune cells present in the tumor microenvironment (TME), affecting the anti-cancer immune response to immunotherapy (Le Bourgeois et al., 2018; Quail and Joyce, 2013).

Metabolic dependencies have been successfully exploited to treat cancer, as illustrated by the efficacy of antifolate drugs such as methotrexate (Walling, 2006). More recent knowledge about cancer cell metabolism has resulted in novel therapeutic targets, such as glutaminase and mutant forms of IDH1/2, currently being evaluated in pre-clinical models and phase I/II clinical trials (Shah and Chen, 2020; Tang et al., 2021). However, adverse effects or lack of effectiveness still hamper the clinical development of most metabolic therapies. A potential reason is that many metabolic targeting drugs are developed based on insights derived from model systems of human cancer, which do not fully reflect the complexities of cancer in humans (Ghaffari et al., 2015). In particular, cell line models lack the immune cells present in the TME (Hynds et al., 2018; Jiang et al., 2016; Vincent and Postovit, 2017) and often require specific metabolic conditions to grow (Ben-David et al., 2018; Hynds et al., 2018).

Evidence is emerging that transcriptional changes play an important role in the metabolic plasticity of cancer cells: gene expression can influence metabolite levels, and metabolic changes can result in altered gene expression (Desvergne et al., 2006; Martin-Martin et al., 2018; Peng et al., 2018). The availability of large numbers of gene expression profiles — from a broad spectrum of cancer types — in the public domain provides a unique opportunity to study metabolic reprogramming in patient-derived cancer tissue.

Almost without exception, these gene expression profiles were generated from complex biopsies that contain tumor cells and cells present in the TME (e.g., immune cells). Accordingly, these profiles represent the average gene expression pattern of all cells present in the biopsy. Therefore, detecting metabolic processes relevant to cancer biology in expression profiles from complex biopsies can be challenging, especially when their transcriptional footprints (TFs) are subtle and concealed by more pronounced TFs from other biological processes or experimental artifacts.

In the present study, we used consensus Independent Component Analyses (c-ICA), a statistical method capable of separating the average gene expression profiles generated from complex biopsies into additive transcriptional components (TCs). This enabled us to detect both the pronounced and more subtle transcriptional footprints of metabolic processes. We performed c-ICA with 32,409 gene expression profiles obtained from the Gene Expression Omnibus (GEO) and The Cancer Genome Atlas (TCGA), as well as 2,085 gene expression profiles obtained from the Cancer Cell Line Encyclopedia (CCLE) and the Genomics of Drug Sensitivity in Cancer portal (GDSC) (Barret et al., 2013; Barretina et al., 2012; Yang et al., 2013). Comprehensive characterization of the TCs with gene set enrichment analysis (GSEA) identified TCs associated with metabolic processes, i.e., metabolic TCs (mTCs). This enabled us to create a metabolic landscape showing the activity of these mTCs in all 34,494 samples. We demonstrate how this landscape (www.themetaboliclandscapeofcancer.com) can be used to explore associations between the metabolic transcriptome and drug sensitivities, patient outcomes, and the composition of immune cells in the TME.

## RESULTS

### A subset of transcriptional components is associated with metabolic processes

Previously, we collected gene expression data from four databases: the Gene Expression Omnibus (GEO dataset, *n* = 21,592), The Cancer Genome Atlas (TCGA dataset, *n* = 10,817), the Cancer Cell Line Encyclopedia (CCLE dataset, *n* = 1,067), and the Genomics of Drug Sensitivity in Cancer (GDSC dataset, *n* = 1,018) (**Figure 1A**), totaling 34,494 samples (Bhattacharya et al., 2020). Overall, 28,200 expression profiles originated from patient-derived complex tissue cancer biopsies, 4,209 from complex tissue biopsies of non-cancerous tissue, and 2,085 from cell lines. The samples in these four databases encompass 89 cancer tissue types and subtypes and 19 non-cancerous tissue types. For GEO and CCLE data sets, the expression profiles were generated with Affymetrix HG-U133 Plus 2.0. Expression profiles within the GDSC dataset were generated with Affymetrix Human Genome U219, and TCGA profiles were generated with RNA sequencing.

**Figure 1.**
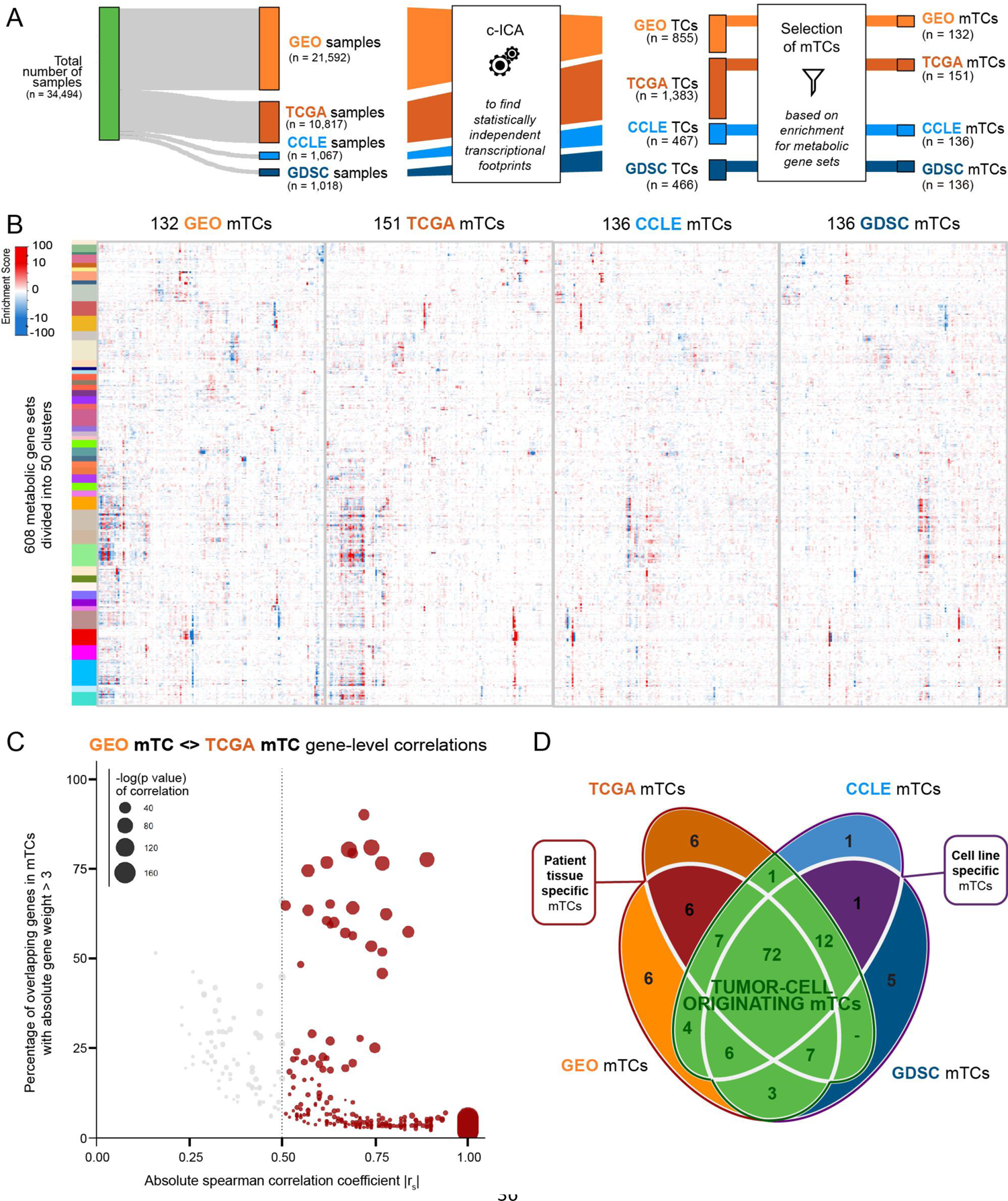
Identification of metabolic transcriptional components (mTCs). **(A)** Workflow for identification of mTCs. Consensus-Independent Component Analysis (c-ICA) applied to identify transcriptional components (TCs). Subsequent systematic selection of TCs enriched for metabolic processes resulted in in 132, 151, 136, and 136 mTCs for the GEO, TCGA, CCLE, and GDSC datasets, respectively. **(B)** Hierarchically clustered heatmaps showing the enrichment of the 608 metabolic gene sets of mTCs identified in GEO, TCGA, CCLE, and GDSC datasets. **(C)** Scatter plot showing absolute spearman correlation coefficients (x-axis), versus the percentage of overlapping top genes (genes with absolute weight >3) between GEO mTCs and TCGA mTCs (y-axis). Only significant pair-wise correlations (with P-values <0.05) are shown. Colored dots show correlations > 0.5, the size of the dots represent the P-value of these spearman correlations. **(D)** Venn diagram quantifying overlap of mTCs between each dataset based on their pair-wise correlations. Two mTCs are counted as shared between datasets, when they have a high absolute spearman correlation (|r_s_|>0.5). Three groups of (shared) mTCs, mentioned in the text, are designated.

Gene expression profiling measures the net expression level of individual genes, thus reflecting the integrated activity of underlying regulatory factors, including experimental, genetic, and non-genetic factors. To gain insight into the number and nature of these regulatory factors and their effects on gene expression levels, i.e., their transcriptional footprints, we previously performed consensus-independent component analysis (c-ICA) on each of the abovementioned four datasets separately (Bhattacharya et al., 2020), resulting in four sets of transcriptional components (TCs). In every TC, each gene has a specific weight. This weight describes how strongly and in which direction the underlying transcriptional regulatory factor influences the expression level of that gene. c-ICA also provides a ‘mixing-matrix’ per dataset, in which each column corresponds to a TC and each row corresponds to a sample. Values in the mixing matrix are interpreted as measurements of the activity of the TCs in an individual sample; we refer to these as ‘activity scores’. Ultimately, the analysis yielded 855, 1383, 466, and 467 TCs for GEO, TCGA, CCLE, and GDSC datasets, respectively (**Figure 1A**).

Gene set enrichment analysis (GSEA) with 608 gene sets that describe metabolic processes was performed to identify TCs enriched for metabolic processes. The gene sets were selected from the gene set collections Biocarta (*n* = 7), the Kyoto Encyclopedia of Genes and Genomes (KEGG, *n* = 64), the Gene Ontology Consortium (GO, *n* = 508), and Reactome (*n* = 29) within the Molecular Signatures DataBase (MSigDB, v6.1; for the systematic selection strategy see **Methods**). We performed consensus clustering on the enrichment scores of the 608 metabolic gene sets to identify potential biological redundancy in the metabolic gene set definitions (**Figure S1**). This resulted in 50 clusters of gene sets, which can be ascribed to different metabolic themes (**Table S1**). Based on these 50 enrichment clusters, 132 (GEO), 151 (TCGA), 136 (CCLE), and 136 (GDSC) mTCs were defined (**Figure 1A** and **B**); see **Methods** for the systematic selection strategy). These mTCs represent the metabolic transcriptional footprints present in our broad set of samples, i.e., patient-derived samples, cancer cell line samples, and non-cancer samples. However, some of the identified mTCs may capture the transcriptional footprints of experimental factors. Therefore, we investigated how much of the variance in activity scores of each mTC could be explained by experimental batches. For GEO mTCs, experimental batches were determined by the provided GSE identifiers (i.e., experiment series identifiers). For TCGA mTCs, experimental batches were determined by the tissue source site of samples (e.g., 2H, Erasmus MC, esophageal carcinoma). We observed that 12/132 GEO mTCs showed a potential putative batch effect with more than 10% explained variance (**Figure S2A**). However, six of the 12 GEO mTCs with a putative batch effect also explained more than 10% of the variance in gene expression of samples belonging to a single tissue subtype (**Figure S2A**). One of the 151 TCGA mTCs showed a putative batch effect with 20.5% explained variance (**Figure S2B**). This mTC, TCGA mTC 43, also showed tissue-specificity for thymoma, a tissue type that is not present in the GEO dataset. These observations might indicate that the mTCs showing a putative batch effect in fact describe tissue-specific biology of tissues that are only present in a single experiment in our dataset.

### Metabolic TCs are robust across different datasets and platforms

Pair-wise comparison of mTCs between datasets, based on gene weights, showed that 91-99% of mTCs per dataset were highly correlated (|*r_s_*| ≥ 0.5, P-value < 0.05 as a threshold) with at least one mTC identified in another dataset (**Figure 1C**, **D** and **Figure S3A-G**). This indicates that most of the mTCs were cross-platform and cross-dataset robust.

Given the selected correlation threshold (|*r_s_*| ≥ 0.5, P-value < 0.05), 72 mTCs could be identified with a highly similar gene weight pattern in all four datasets (**Figure 1D**). Thus, these mTCs capture a transcriptional footprint that is very similar in both patient-derived complex biopsies and cell lines. As cell lines lack a TME, these 72 mTCs were considered to capture metabolic processes that reflect tumor cell characteristics. Six GEO mTCs were identified that were highly correlated with TCGA mTCs, but not highly correlated with any CCLE or GDSC mTC (**Figure 1D**). These mTCs, therefore, might capture transcriptional footprints that are specific for complex biopsies obtained from patient-derived cancer tissue and may originate from the TME or capture a transcriptional footprint from tissue only present in the GEO and TCGA datasets. One pair of mTCs was identified with a gene weight pattern that was highly similar in CCLE and GDSC datasets only, capturing a metabolic transcriptional footprint that could only be found in cell line models (**Figure 1D**).

### Metabolic TCs identify new genes potentially involved in metabolic processes

Among the ‘top’ genes in every mTC —defined as the genes with an absolute weight > 3 in an mTC — many genes were member of the 608 metabolic gene sets (**Figure 2A** and **2B, Figure S4**).

**Figure 2.**
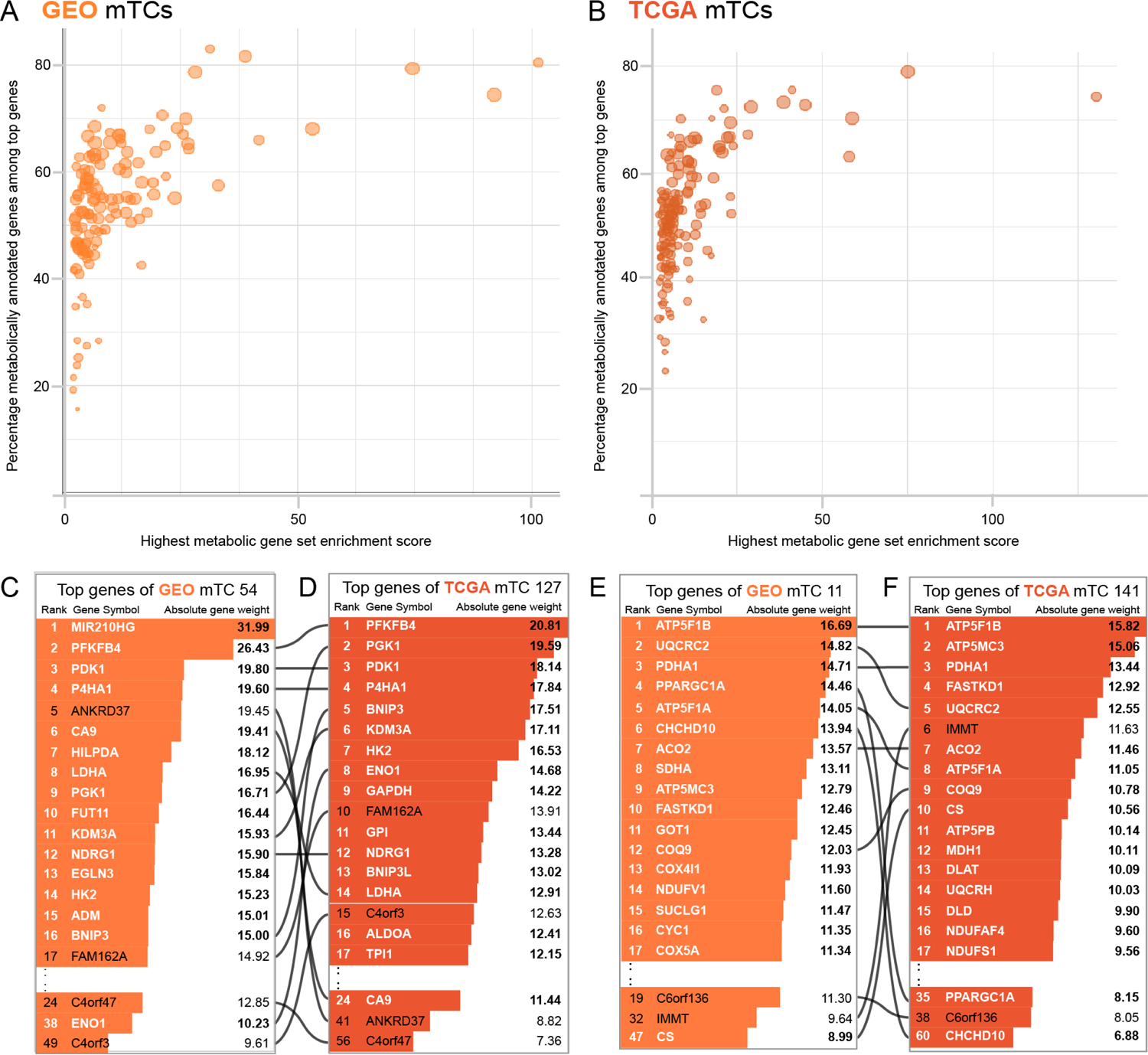
Metabolic TCs identify new genes potentially involved in metabolic processes. **(A-B)** Scatterplots showing the highest metabolic gene set enrichment score for every GEO (A) and TCGA (B) mTC (x-axis) versus the percentage of metabolically annotated genes among the top genes (genes with absolute weight >3) in those mTCs. Size of dots correspond to the absolute amount of metabolically annotated genes in the corresponding mTC. **(C-D)** Top genes in GEO mTC 54 and TCGA mTC 127. Text colored white shows genes that are a member of at least one of the 608 defined metabolic gene sets. Lines signify genes that are top genes in both GEO and TCGA mTCs. **(E-F)** Top genes in GEO mTC 11 and TCGA mTC 141.

However, even for the mTCs with the absolute highest gene set enrichment scores for a metabolic gene set, at least 20% of top genes were not members of any of the metabolic gene sets. Because these genes were nevertheless part of an mTC, they may be potentially involved in the metabolic processes that showed enrichment.

For example, two strongly correlated mTCs, GEO mTC 54 and TCGA mTC 127 (|*r_s_*|= 0.77), both showed enrichment for glycolysis and the metabolic process of ADP (**Figure 2C** and **2D, Table S1**). GEO mTC 54 contained 262 top genes, of which 155 (59.1%) were also among the top genes in TCGA mTC 127. Both mTCs contained multiple top genes that are known targets of the HIF-1 complex, and genes previously found to be part of a hypoxic signature (Benita et al., 2009; Ye et al., 2018). Several top genes of both GEO mTC 54 and TCGA mTC 127 (e.g., *FAM162A*, *C4orf3*, *C4orf47*, and *ANKRD37*) are currently not a member of any of the 608 metabolic gene sets. However, these data suggest that these four genes are involved in glycolysis and are possibly hypoxia related. Indeed, several studies have indicated that at least *FAM162A* and *ANKRD37* are regulated by the transcription factor HIF-1α (Copple et al., 2012; Sørensen et al., 2015).

As a second example, we investigated two highly correlated mTCs, GEO mTC 11 and TCGA mTC 141 (*r_s_* = 0.68), which showed enrichment for mitochondrial metabolic processes such as oxidative phosphorylation and the TCA cycle (**Figure 2E** and **2F, Table S1**). GEO mTC 11 contained 427 top genes, of which 270 (63.2%) were among the top genes in TCGA mTC 141. In these two mTCs, *C6orf136* and *IMMT* are top genes currently not assigned to any of the 608 metabolic gene sets. *C6orf136* and *IMMT* were previously identified in functional mitochondria’ proteome profiles (Lefort et al., 2009). These results suggest that mTCs could assign metabolic functions to genes currently not members of known gene sets describing metabolic processes.

### Clustering sample activity scores of mTCs reveal multiple metabolic subtypes

To investigate the heterogeneity of the metabolic transcriptome in a broad range of cancer subtypes, we hierarchically clustered the mixing matrix provided by consensus-ICA that contains the activity score of mTCs in every sample (**Figure 3A, 3B** and **S5A, S5B**). We selected the cutoff heights of the resulting dendrograms so that every cluster – referred to as metabolic subtype – contained at least 50 samples (**Figure S5C** and **S5D**). This clustering analysis divided the 21,592 GEO samples into 67 metabolic subtypes with a median of 276 samples per subtype (range 54-1,252) and the 10,817 TCGA samples into 58 metabolic subtypes with a median of 167 samples per subtype (range 52-536). For an overview of the metabolic subtypes and their sample composition, see **Figure S6, S7**, and **Table S2**. Two types of patterns emerged.

**Figure 3.**
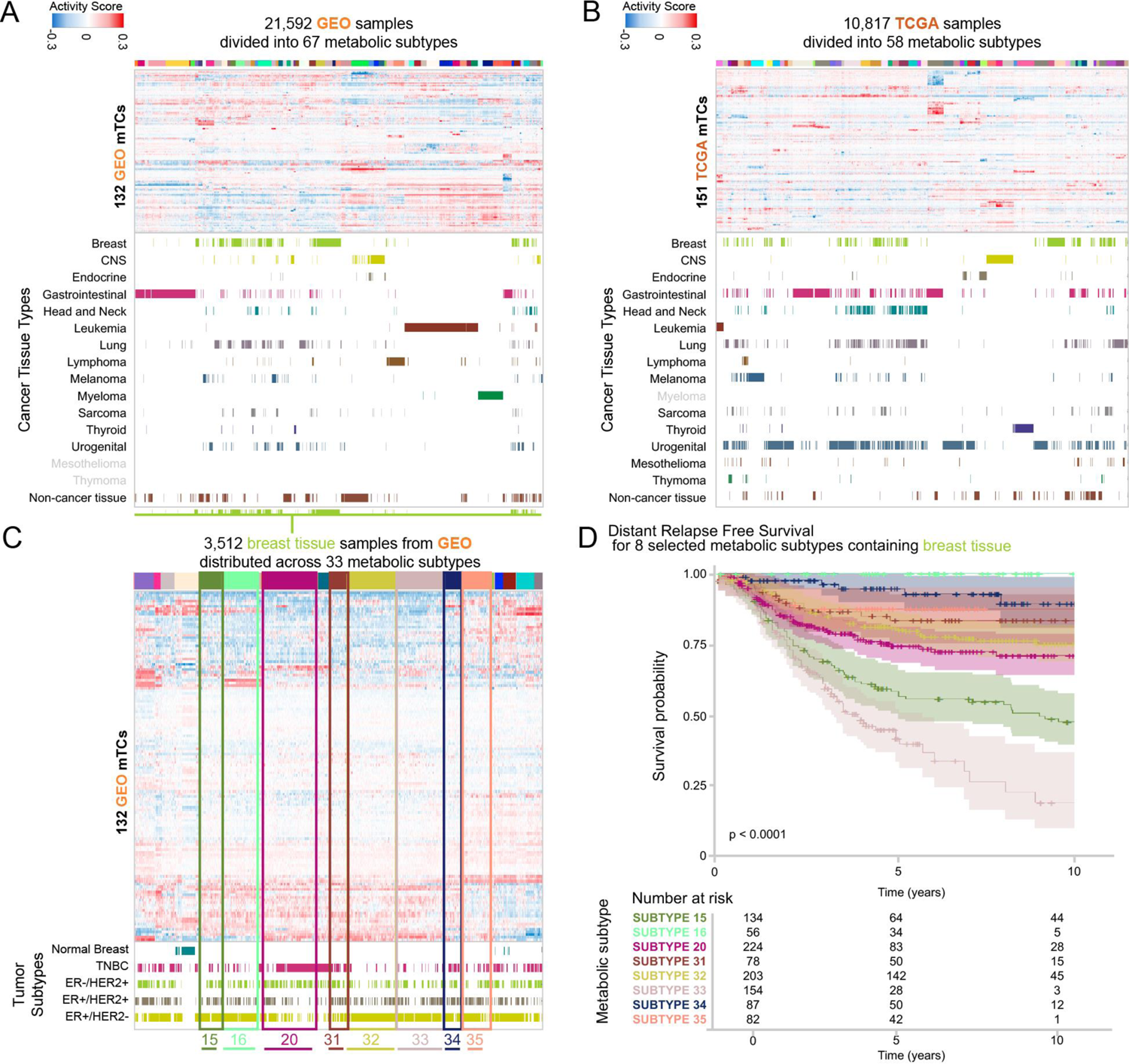
Clustering activity scores of mTCs reveal multiple metabolic subtypes. **(A)** 21,592 GEO samples were hierarchically clustered based on mTC activity scores and divided into 67 metabolic subtypes. **(B)** 10,817 TCGA samples were hierarchically clustered based on mTC activity scores and divided into 58 metabolic subtypes. **(C)** Metabolic landscape of the subset of breast tissue samples in the GEO dataset. Subtypes with DFS data were selected for survival analysis are highlighted. Grey labels designate tissue types that are present in other datasets, but are not present in the given dataset. **(D)** Distant relapse-free survival of breast cancer patients in the GEO dataset. Patient-derived samples were stratified per metabolic subtype. Kaplan Meier curves are shown with a confidence interval of 0.95.

The first pattern consisted of tumor types with samples that belong to one dominant metabolic subtype. For example, 102/133 (76.7%) of thyroid cancer samples in the GEO dataset fell into one metabolic subtype (subtype 27, **Figure S6, Table S2**). Similarly, 446/509 (87.6%) of thyroid cancer samples in the TCGA dataset fell into metabolic subtype 43 (**Figure S7, Table S2**). In line with the biology of thyroid tissue, both GEO metabolic subtype 27 and TCGA metabolic subtype 43 were characterized by high activity scores of mTCs enriched for thyroid hormone metabolism (GEO mTC 64 and TCGA mTC 87; **Table S1**).

The second pattern consisted of several tumor types that were not characterized by a few dominant metabolic subtypes. Instead, their samples were divided across multiple metabolic subtypes. For example, the 3,512 breast cancer samples in the GEO dataset were divided across 33 metabolic subtypes (**Figure 3C**). These metabolic subtypes did not follow the breast cancer classification based on ER and HER2 receptor status (**Figure 3C** and **Table S2**). In line with this observation in the GEO dataset, the 1,100 breast cancer samples in the TCGA dataset were also scattered across 29 metabolic subtypes.

Several metabolic subtypes likewise contained samples from multiple tumor types. For example, GEO metabolic subtype 22 contained samples from 25 tumor types, including 42 ovarian cancer samples (22% of all ovarian cancer), 33 synovial sarcoma samples (97% of all synovial sarcoma), and 15 Ewing’s sarcoma samples (58% of all Ewing’s sarcoma; **Figure S6** and **Table S2**). GEO mTC 111 had the highest absolute median activity score in GEO metabolic subtype 22 (**Table S2**). This mTC showed enrichment for the metabolism of nicotinamide adenine dinucleotide phosphate (NADP) and genes involved in the activation of an innate immune response (**Table S1**). These results show that the classification of samples based on metabolic subtype yields different patterns than current classification systems, such as histotype or receptor status in breast cancer.

### Metabolic subtypes are associated with distant relapse-free survival in breast cancer

We then investigated if metabolic subtypes could have clinical relevance. We had previously collected distant relapse-free survival (DRFS) data for 1,207 breast cancer samples (Bense et al., 2017). As mentioned earlier, breast cancer samples in the GEO dataset were divided across 33 of the 67 metabolic subtypes. Of these 33 subtypes, eight contained > 50 breast cancer samples with data available for DRFS: subtypes 15, 16, 20, 31, 32, 33, 34, and 35. We found that patients from breast cancer samples assigned to metabolic subtypes 16 and 33 showed the best and worst DRFS, respectively (P-value = 1.08·10^-23^, Log-Rank test; **Figure 3D**). Distributions of standard prognostic factors within these eight metabolic subtypes are presented in **Table S3**. These results show that metabolic subtypes are associated with disease outcomes in breast cancer.

### The activity of mTCs is associated with drug sensitivity

The CCLE and GDSC databases contain the sensitivities of cell lines to a large panel of drugs expressed as IC_50_ values. With a threshold of |r_s_|>0.2, we observed associations between the activity scores of 61 CCLE mTCs, 90 GDSC mTCs, and the IC50 values of 238 drugs (**Table S4**).

For example, in the GDSC dataset, an increase in activity score of GDSC mTC 3 was associated with a decrease in IC_50_ value of (i.e., increased sensitivity to) nutlin-3a (|*r_s_*|= 0.42; **Figure 4A** and **4B**). Nutlin-3a targets the p53 pathway through inhibition of MDM2. In line with this, GDSC mTC 3 showed strong enrichment for genes involved in the p53 pathway, with *MDM2* ranked as the second gene (**Table S1**). GDSC mTC 3 was strongly correlated with CCLE mTC 4 (|*r_s_*|= 0.84), GEO mTC 57 (|*r_s_*|= 0.79), and TCGA mTC 110 (|*r_s_*|= 0.74) (**Figure 4D**), suggesting that this mTC was captured in cell line datasets as well as in the two patient-derived datasets. Indeed, an increase in activity score of CCLE mTC 4 was associated with a decrease in IC_50_ value of nutlin-3a as well (|*r_s_*|= 0.25; **Figure 4E**). Cell lines with wildtype *TP53* had a higher activity score of GDSC mTC 3 (**Figure 4C**). Also, cell lines with wildtype *TP53* had a higher activity score of CCLE mTC 4 (**Figure 4F**).

**Figure 4.**
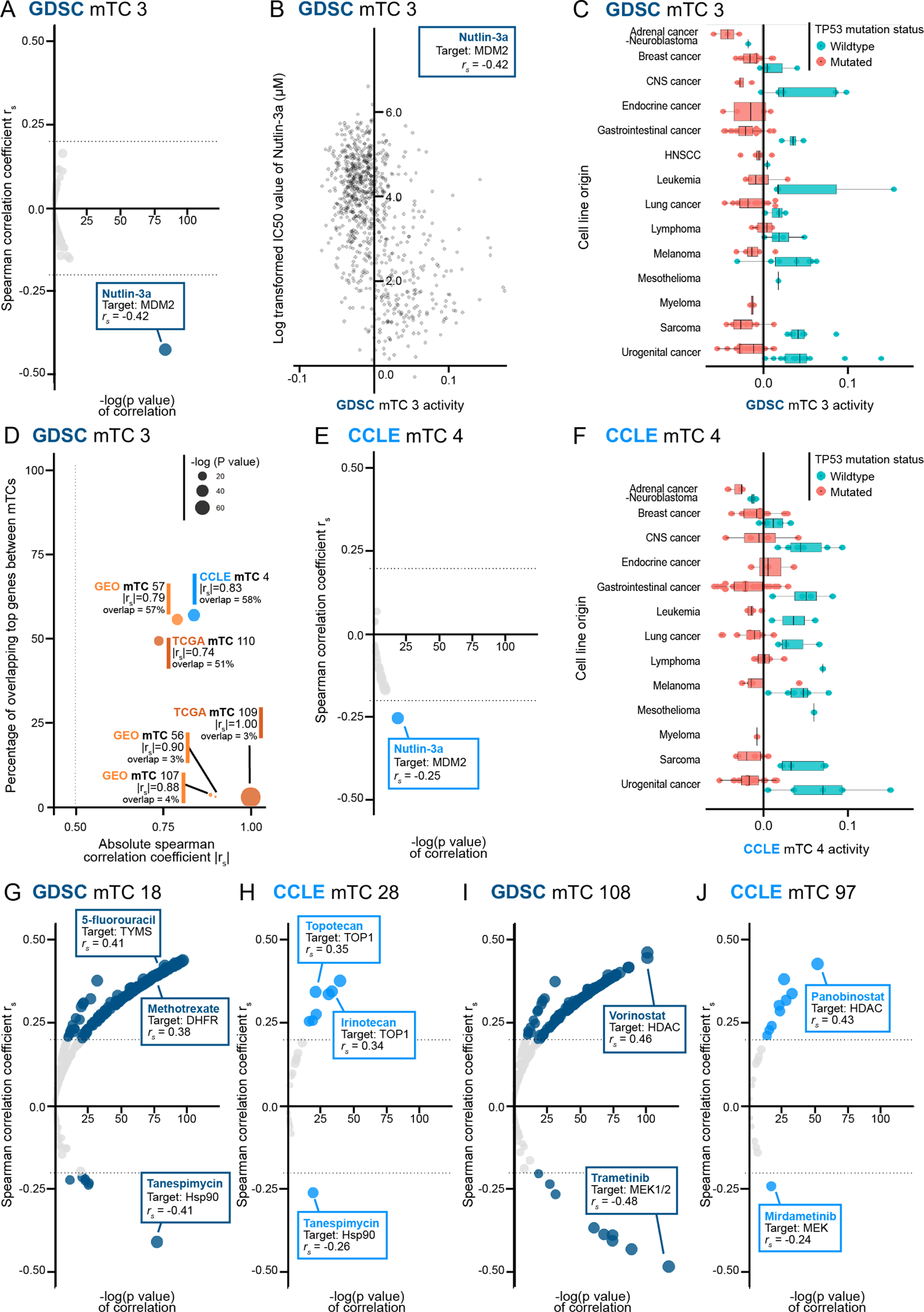
Associations between mTCs and drug sensitivity for selected examples. **(A)** Spearman correlations between drug IC50 values and the activity of GDSC mTC 3 **(B)** Scatter plot showing the association between the (log-transformed) IC50 value of Nutlin-3a and activity of GDSC mTC 3 in samples. **(C)** Box plot of activity of GDSC mTC 3 across cell lines, colored for their *TP53* mutation status. **(D)** Pair-wise correlations between GDSC mTC 3 and mTCs from GEO, TCGA and CCLE datasets. Every dot corresponds to an mTC with a correlation to GDSC mTC 3 ≥ 0.5. Dot sizes correspond to the P-value of the spearman correlation coefficient; the y-axis gives the percentage of overlapping top genes between the two mTCs involved in the correlation. **(E)** Spearman correlations between drug IC50 values and the activity of CCLE mTC 4 **(F)** Box plot of activity of CCLE mTC 4 across cell lines, colored for their *TP53* mutation status. **(G-J)** Spearman correlations between drug IC50 values and the activity of GDSC mTC 18, CCLE mTC 28, GDSC mTC 108 and CCLE mTC 97.

In another example, the activity score of GDSC mTC 18 was found to be associated with the IC50 values of 142 drugs (|*r_s_*| range 0.20 – 0.44; **Figure 4G**). An increase in activity score of GDSC mTC 18 in a sample was associated with a higher IC_50_ value (i.e., increased resistance) for 135 of these drugs, including the widely used DNA synthesis-inhibiting antimetabolites 5-fluorouracil (|*r_s_*| = 0.41) and methotrexate (|*r_s_*| = 0.38). GDSC mTC 18 was strongly correlated with CCLE mTC 28 (|*r_s_*|= 0.84), GEO mTC 35 (|*r_s_*|= 0.59), and TCGA mTC 58 (|*r_s_*|= 0.55), indicating that this mTC is also captured in both cell line datasets and the two patient-derived datasets. In line with this, CCLE mTC 28 was associated with a higher IC_50_ value (i.e., increased resistance) for 7 drugs including topoisomerase inhibitors topotecan (|*r_s_*| = 0.35) and irinotecan (|*r_s_*| = 0.34) (**Figure 4H**). All four of the highly correlated mTCs were enriched for genes involved in glutathione metabolism, cellular ketones and xenobiotics, and drug detoxification (**Table S1**). Specifically, genes belonging to the aldo-keto reductase family 1 (AKR1) were among the top genes in these mTCs. Previous studies have reported a role for the glutathione system in resistance to irinotecan and 5-fluorouracil (Goto et al., 2002), and specifically the AKR1 family in resistance to e.g. methotrexate and irinotecan (Heibein et al., 2012; Matsunaga et al., 2020; Selga et al., 2008). In contrast, we observed that an increased activity score of GDSC mTC 18 was associated with a decrease in IC_50_ value (i.e., increased sensitivity) for only seven drugs (|*r_s_*| range 0.20-0.41; **Figure 4G**). The drug with the highest negative correlation was tanespimycin (17-AAG), an Hsp90 inhibitor (|*r_s_*| = 0.41). An increased activity score of CCLE mTC 28 was associated with decrease in IC_50_ value for tanespimycin as well (|*r_s_*| = 0.26; **Figure 4H**). A direct link between the functions of glutathione and Hsp90 in oxidative stress has been suggested, as well as a relationship between tanespimycin sensitivity and *NQO1* expression, a gene coding for an enzyme reducing quinones to hydroquinones that is involved in detoxification pathways (Gaspar et al., 2009; Kim et al., 2015). In line with these findings, we found that the *NQO1* gene is present near the top of GDSC mTC 18, CCLE mTC 28, GEO mTC 35, and TCGA mTC 58.

As a final example, increased activity of GDSC mTC 108 was associated with a lower IC_50_ value (i.e., increased sensitivity) to the MEK inhibitor trametinib (|*r_s_*| = 0.48) and a higher IC_50_ value (i.e., increased resistance) to the histone deacetylase inhibitor vorinostat (|r_s_| = 0.46; **Figure 4I** and **Table S4**). GDSC mTC 108 was correlated with CCLE mTC 97 (|r_s_|= 0.32). Consistent with the observation for GDSC mTC 108, we found that increased activity of CCLE mTC 97 was associated with a lower IC_50_ value (i.e., increased sensitivity) to the MEK inhibitor mirdametinib (|*r_s_*| = 0.24) and a higher IC_50_ value (i.e., increased resistance) to the histone deacetylase inhibitor panobinostat (|r_s_| = 0.43; **Figure 4J** and **Table S4**). This contrasting sensitivity for MEK and histone deacetylase inhibition is in line with data from a study that used *BRAF*-mutated melanoma cell lines. The authors showed that cell lines with acquired resistance to MEK inhibitors subsequently became sensitive to treatment with the histone deacetylase inhibitor vorinostat (Wang et al., 2018). They concluded that the MEK-inhibitor resistance mechanism results from the activation (or reactivation) of MAPK cascades (Wagle et al., 2014). These findings are in line with our observation that both GDSC mTC 108 and CCLE mTC 97 were enriched for genes involved in the negative regulation of the MAPK cascade (**Table S1**). These examples demonstrate how mTCs can capture cross-dataset robust metabolic transcriptional footprints relevant for drug response.

### The activity of mTCs is associated with the immune composition of the tumor microenvironment

We determined the association between the activity of mTCs and the immune composition of the TME (**Table S5**; see **Methods** for details). The immune composition for all samples in the GEO and TCGA dataset was determined by inferring fractions of 22 immune cell types using the CIBERSORT algorithm (Chen et al., 2018). We observed that the mTCs that were correlated with immune cell fractions could be divided into two groups. The first group included mTCs that were only identified in the patient-derived datasets. The second group contained mTCs that were identified in both the patient-derived and the cell line datasets.

For example, the activity score of GEO mTC 123 was associated with estimated fractions of CD8+ T cells (|*r_s_*| = 0.40), γδ T cells (|*r_s_*| = 0.36), activated CD4 memory T cells (|*r_s_*| = 0.34), and regulatory T cells (|*r_s_*| = 0.32, **Figure 5A**). Belonging to the group of mTCs only identified in the patient-derived datasets, GEO mTC 123 was correlated highly with only TCGA mTC 34 (|*r_s_*|= 0.28).

**Figure 5.**
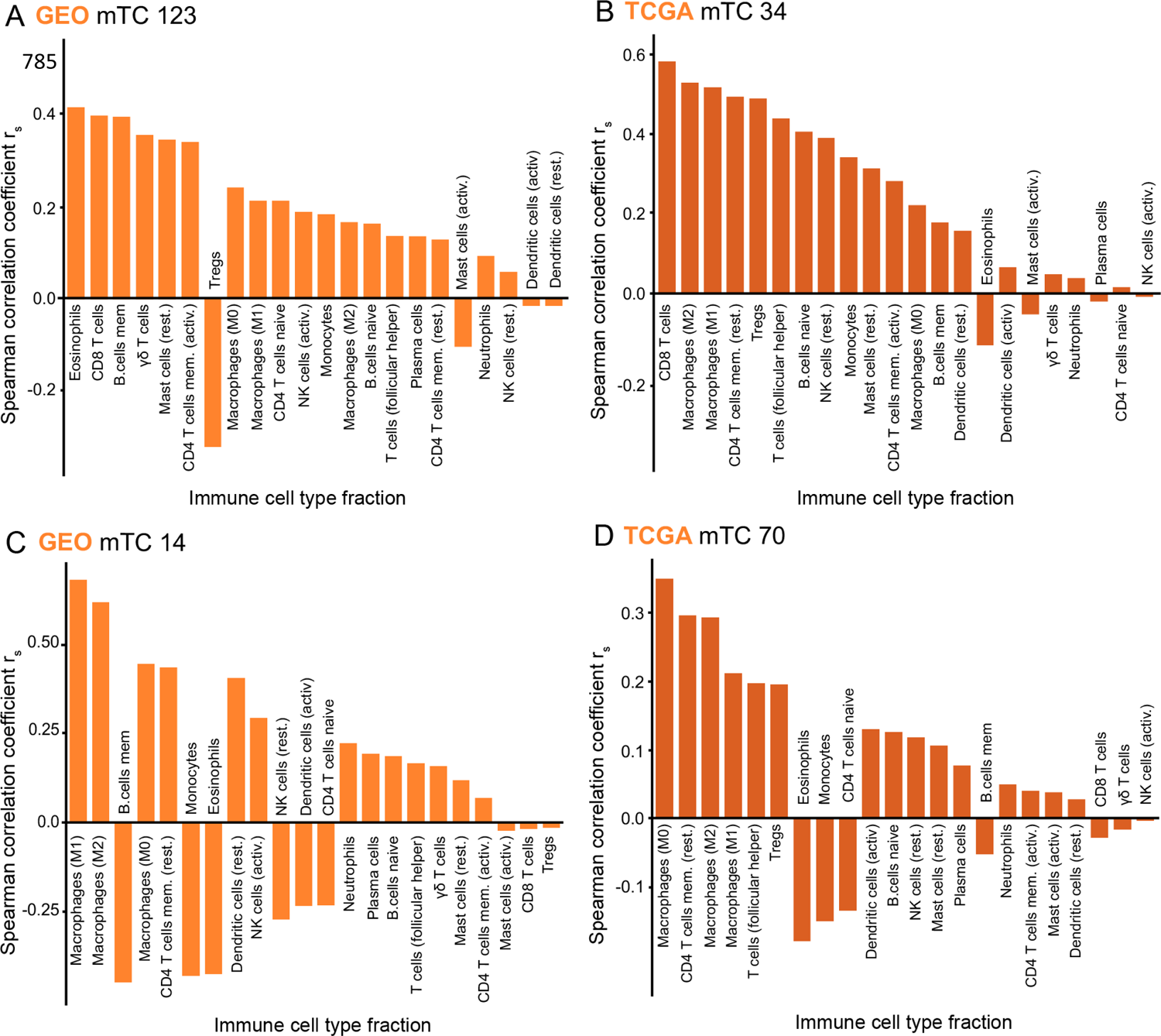
Associations between mTCs and the composition of the immune tumor microenvironment for selected examples. **(A-B)** Spearman correlations between CIBERSORT estimated immune cell fractions and the activity of GEO mTC 123 and TCGA mTC 34. **(C-D)** Spearman correlations between CIBERSORT estimated immune cell fractions and the activity of GEO mTC 14 and TCGA mTC 70.

In line with this, the activity score of TCGA mTC 34 was also associated with CD8+ T cell fractions (|*r_s_*| = 0.58, **Figure 5B**). Both GEO mTC 123 and TCGA mTC 34 showed enrichment for genes involved in immunological processes such as leukocyte activation and cytokine metabolism and metabolic processes such as phosphatidylinositol and phospholipid metabolism (**Table S1**). The fact that both GEO mTC 123 and TCGA mTC 34 have no high correlation to mTCs in the cell line datasets suggests that they indeed capture transcriptional activity from non-cancerous cells in the immune TME.

GEO mTC 14 is illustrative of the second group of mTCs correlated with immune cell fractions and identified in both the patient-derived and the cell line datasets. The activity scores of GEO mTC 14 were correlated with the fractions of M1 macrophages (|*r_s_*|= 0.65) and M2 macrophages (|*r*_s_|= 0.59; **Figure 5C**). GEO mTC 14 was correlated with TCGA mTC 70 (|*r_s_*|= 0.44) and with CCLE mTC 124 (|*r_s_*|= 0.47), and GDSC mTC 33 (|*r_s_*|= 0.33). All four mTCs were enriched for genes involved in the metabolism of extracellular macromolecules (**Tables S1**). Genes coding for several types of collagens were among the top-ranked in these mTCs. This is in line with previous reports indicating that macrophages can function as collagen-producing cells in the TME (Schnoor et al., 2008; Vaage and Harlos, 1991). GEO mTC 14 and TCGA mTC 70 showed a high activity score in subsets of breast cancers, lung cancers, and sarcomas (**Figure S8A** and **S8B**). A negative activity score of GEO mTC 14 and TCGA mTC 70 was observed in a subset of hematological cancers and hematological cancer cell lines in both GDSC and CCLE mTCs. Because these mTCs were present in both patient data sets and cell line datasets, this indicates that the captured metabolic processes reflect tumor cell characteristics, which are associated with the fraction of macrophages present in the immune TME.

By correlating inferred immune cell fractions of samples with the activity scores of mTCs in samples, the relationship between the metabolic transcriptome and the various components of the immune TME could be assessed.

## DISCUSSION

In the present study, we used consensus-Independent Component Analysis (c-ICA) in combination with Gene Set Enrichment Analysis (GSEA) to identify a broad set of robust metabolic Transcriptional Components (mTCs). With these mTCs, the transcriptional metabolic landscape was defined in patient-derived cancer tissue, cancer cell lines, and non-cancer samples. We also showed how this metabolic landscape could be used to generate hypotheses by exploring associations between metabolic processes and drug sensitivities, patient outcomes, and the composition of the immune tumor microenvironment.

We used the wealth of publicly available pan-cancer transcriptomic data to study human metabolism on a large scale. Previous work used either single-cell sequencing or bulk cell transcriptomic profiles to study metabolism in specific cancer types (Hakimi et al., 2016; Xiao et al., 2019), or pan-cancer, but based on a single platform (Cubuk et al., 2018; Rosario et al., 2018). Our present study differs from this previous work in two essential aspects. Firstly, we used c-ICA to segregate the average expression patterns of complex biopsies into statistically independent components (Biton et al., 2014; Kong et al., 2008). Previous studies investigated average gene expression profiles of complex biopsies and can therefore only distinguish the gene expression signature and regulation of more pronounced metabolic processes. With c-ICA it is possible to identify statistically independent regulatory factors and their transcriptional footprints and distinguish both pronounced from more subtle metabolic processes. This enabled us to determine the association of both pronounced and subtle metabolic processes with, e.g., patient outcome and the composition of the TME in a complex biopsy. Secondly, the present study is the most extensive transcriptional analysis of metabolism and the first that integrated patient-derived data from GEO and TCGA with cell line data from CCLE and GDSC. The samples in these four datasets were obtained from a multitude of independently constructed, publicly available cohorts, and the expression profiles were generated using different technologies (microarray or RNA-sequencing). This integrated dataset enabled us to demonstrate that most of the identified mTCs were robust and independent from dataset-specific and platform-specific characteristics. The observed overlap, or lack of overlap, between patient-derived and cell line-derived mTCs can help researchers understand how metabolic genes and pathways identified in cell lines can be translated to a patient tissue context and vice versa.

Furthermore, we hypothesize that metabolic processes identified only in patient-derived samples and not in cell line samples are more likely to originate from cells in the tumor microenvironment. These microenvironment-specific metabolic processes will not be captured by mTCs in cell line datasets. This is because bulk expression profiles of cancer cell line samples do not harbor transcriptional footprints associated with non-cancerous cells.

The metabolic landscape enabled us to classify samples based on the transcriptional activity of metabolic processes, resulting in metabolic subtypes. However, this metabolic classification was often not in full alignment with current classification systems based on aspects such as histotype. We demonstrated that metabolic subtypes were associated with disease outcomes for breast cancer, emphasizing the relevance of metabolic pathway-based classification in cancer.

The heterogeneity (metabolic and otherwise) within and between cancer types is well recognized, and alternative subtyping based on metabolite profiling and the metabolic transcriptome have been proposed before (Reznik et al., 2018; Rosario et al., 2018; Tang et al., 2014). More specifically, clinically significant metabolism-based classifications have been proposed in breast cancer (Cappelletti et al., 2017; Serrano-Carbajal et al., 2020; Wang et al., 2019). The most active mTCs in a metabolic subtype relevant to disease outcome could thus be used to generate new hypotheses for treatment targets. Additionally, the association between the activity of mTCs and drug sensitivity could help to design these future therapeutic strategies.

Metabolic heterogeneity and plasticity are not limited to cancer cells but are also applicable to the immune cells present in the tumor micro-environment. Immune cells undergo metabolic changes when activated, and their metabolic status can overlap with the metabolic state of cancer cells (Andrejeva and Rathmell, 2017). For example, the Warburg effect is classically seen as an example of a metabolic transformation in cancer cells. However, it is also observed in activated T cells (Bantug et al., 2018; Patel and Powell, 2017; Wang and Green, 2012). In the context of metabolism, this complex interplay between cancer cells and immune cells present in the micro-environment gives a new dimension to the use of drugs that target metabolic processes (O’Sullivan et al., 2019; Patel et al., 2019). For instance, inhibiting glutamine metabolism has been shown to inhibit tumor growth and increase the sensitivity of triple-negative breast cancers to immune checkpoint blockade (Oh et al., 2020), and reducing oxidative stress has been shown to prevent the generation of tumor-associated macrophages (Zhang et al., 2013). Furthermore, modulating metabolism in T cells from glycolytic to an OXPHOS-weighted profile has been shown to improve CAR T cell immunotherapy (Fraietta et al., 2018; O’Sullivan and Pearce, 2015; Sukumar et al., 2017). Our transcriptional metabolic landscape can contribute to knowledge on immunometabolism and, combined with the association of mTCs with drug sensitivity, also contribute to the formulation of new hypotheses on how to metabolically engage the tumor and its immune microenvironment, thus improving the response to immunotherapy.

Further research to gain an even more comprehensive understanding of metabolism in patient-derived cancer samples should ideally integrate genomics, transcriptomics, proteomics, and metabolomics to capture the complexity of metabolic processes within cancer cells (Buescher and Driggers, 2016). Recent initiatives are the Recon1, Edinburgh Human Metabolic Network (EHMN), and Human1 projects (Brunk et al., 2018; Ma et al., 2007; Robinson et al., 2020). However, challenges for these initiatives lie in the limited set of samples for which these high-dimensional multi-omics features are available and the use of predominantly cell line samples. Paired datasets on a large scale are needed to unleash the full potential of such an integrated approach.

To facilitate the use of our transcriptional metabolic landscape, we have provided access to all data via a web portal (www.themetaboliclandscapeofcancer.com). In this portal, users can explore genes, metabolic processes, and tissue types of interest. We invite researchers and clinicians to use this portal as a guide to the metabolic transcriptome in cancer or as a starting point for further research into cancer metabolism.

## MATERIALS AND METHODS

### Resource availability

Further information and requests for resources should be directed to the Lead Contact, Rudolf S.N. Fehrmann.

## Data and code availability

Data can be explored at http://themetaboliclandscapeofcancer.com. Code is available at github.com/MetabolicLandscape/

### Data acquisition

A detailed description of the data acquisition of the four datasets has been described previously (Bhattacharya et al., 2020). In short, the GEO dataset contained microarray expression data generated with Affymetrix HG-U133 Plus 2.0 (accession number GPL570). A two-step search strategy was applied to select healthy or cancer tissue samples – automatic filtering on keywords followed by manual curation. Samples from cell lines, cultured human biopsies, and animal-derived tissue were excluded. The TCGA dataset contained the preprocessed and normalized level 3 RNA-seq (version 2) data for 34 cancer datasets available at the Broad GDAC Firehose portal (https://gdac.broadinstitute.org/). The profiles in the CCLE dataset were generated with Affymetrix HG-U133 Plus 2.0. The CCLE project conducted a detailed genetic characterization of a large panel of human cancer cell lines. Expression data within the CCLE project was generated with Affymetrix HG-U133 Plus 2.0. The GDSC dataset contained expression data generated with Affymetrix HG-U219. The GDSC project aims to identify molecular features of cancer that predict response to anti-cancer drugs.

### Preprocessing, normalization, and quality control

A more detailed description has been provided previously (Bhattacharya et al., 2020). In short, preprocessing and aggregation of raw expression data (CEL files) within the GEO dataset, CCLE dataset, and GDSC dataset was performed according to the robust multi-array average algorithm RMAExpress (version 1.1.0). Quality control was performed on the GEO dataset, CCLE dataset, and GDSC dataset separately with principal component analysis (PCA). Duplicate CEL files were removed by generating a message-digest algorithm 5 (MD5) hash for each CEL file. The expression levels for each probeset (in the GEO dataset, CCLE dataset, and GDSC dataset) or gene (in the TCGA dataset) were standardized to a mean of zero and variance of one to remove probeset-specific or gene-specific variability in the datasets.

### Consensus independent component analysis

We used consensus independent component analysis (c-ICA) to segregate the average gene expression patterns of complex biopsies into statistically independent transcriptomic components. The input gene expression dataset was preprocessed using whitening transformation, making all profiles uncorrelated and giving them a variance of one. Next, ICA was performed on the whitened dataset using the FastICA algorithm, resulting in the extraction of estimated sources (ESs) and a mixing matrix (MM). The number of principal components which captured 90% of the variance seen in the whitened dataset was chosen as the number of ESs to extract. Each ES contains all genes with a specific weight. This weight represents the direction and magnitude of the influence of an underlying transcriptional regulatory process on that gene expression level. The MM contains the coefficients of ESs in each sample, representing the activity of an ES in the corresponding sample. We performed 25 ICA runs with different random initialization weight factors to assess the robustness of the ESs and exclude ICA results derived from convergence at local solutions. ESs extracted from these runs were clustered together if the absolute value of the Pearson correlation between them was > 0.9. We calculated consensus transcriptional components (TCs) by taking the mean vector of weights in the co-clustering ESs. We considered a consensus TC robust when clustering included individual TCs from > 50% of the runs. The consensus TCs, in combination with the original input expression profiles, were used to obtain the consensus mixing matrix (MM) with the individual activity scores of the consensus TCs in each sample via matrix inversion.

### Identification of transcriptional components enriched for metabolic processes

First, we selected gene sets defining metabolic processes from five gene set collections obtained from the Molecular Signatures Database (MSigDb version 6.1); BioCarta, Gene Ontology – Biological Process (GO-BP), Gene Ontology – Molecular Function (GO-MF), KEGG, and Reactome. From BioCarta, gene sets were selected manually based on their title. Selected gene sets described metabolic pathways or regulatory pathways regulated by metabolic processes. From GO-BP, all gene sets were selected that contained the motif ‘METABOLIC_PROCESS’ in the title. Also, gene sets that included the name of a metabolite or class of metabolites combined with the motif’_TRANSPORT’ in the title were selected.

Furthermore, gene sets not containing these title motifs but associated with (cancer) metabolism were manually selected based on metabolic pathway names. From GO-MF, all gene sets were selected that included the name of a metabolite or class of metabolites combined with the motif ‘_ACTIVITY’ or ‘_BINDING’ in the title. From KEGG, all gene set containing the motif “METABOLISM” or “BIOSYNTHESIS” in the title in combination with the name of a known metabolic route was selected. Furthermore, gene sets concerning metabolism-related regulatory pathways were chosen based on their titles. From Reactome, all gene set that falls within the hierarchy of the “Metabolism”-pathways were selected (see reactome.org/PathwayBrowser). The metabolism of Abacavir was not included. A complete list of all metabolic gene sets selected is presented in **Table S1**.

To identify transcriptional components enriched for metabolic processes, gene set enrichment analysis (GSEA) was performed using the selected metabolic gene sets. Enrichment of each metabolic gene set was tested according to the two-sample Welch’s t-test for unequal variance between the metabolic set of genes under investigation versus the set of genes that was not under investigation. To compare gene sets of different sizes, we transformed Welch’s t statistic to a Z-score.

A biological process can be captured by multiple gene sets in several gene set collections. Therefore, it is possible that within the selection of 608 gene sets, multiple gene sets describe the same metabolic process. These will then show a similar pattern in gene set enrichment scores of transcriptional components. To reduce this redundancy, consensus clustering was performed gene set-wise on the GSEA data for the GEO, TCGA, CCLE, and GDSC datasets. Consensus clustering was performed using the ConsensusClusterPlus-package (v1.51.1) within R, using the default hierarchical clustering algorithm and Pearson correlation distance, a maximum amount of clusters (maxK) of 150, 2000 resamplings (reps), with 80% row and 80% column resampling (pFeature and pItem, respectively). The optimal number of clusters (*k)* was determined as the *k* at which the relative change in area under the CDF curve was minimized (<0.01). This resulted in a *k* of 50 clusters (**Figure S1**).

The 50 clusters of gene sets were subsequently used to select transcriptional components based on their enrichment for metabolic processes. Per gene set cluster, the three TCs with the highest absolute enrichment score for any gene set in that cluster were selected. In addition to this, the three TCs with the highest absolute mean enrichment score for all gene sets in that cluster were selected. The selected TCs were then referred to as metabolic Transcriptional Components (mTCs). In the end, four different sets of mTCs were identified (GEO mTCs, TCGA mTCs, CCLE mTCs, GDSC mTCs)

### Approximation of batch effects and tissue specificity of mTCs

First, the explained variance of every component from the perspective of a sample (as a percentage) was estimated using the squares of the mixing matrix weights of a sample divided by the sum of the squares. This percentage explained variance matrix for samples was then summarized into a mean explained variance for studies by summarizing samples belonging to the same study (through the annotated GEO series accession number or TCGA tissue source site code). In the figures, only the highest explained variance available for any study is given. Similarly, tissue specificity was approximated by calculating the mean explained variance for tissue types by summarizing samples belonging to the same tissue subtype.

### Pair-wise gene-level correlations of mTCs between datasets

To correlate two mTCs of different datasets, the subset of genes with an absolute weight higher than 3 in two mTCs was selected. Then, the overlap between these two sets of top genes was determined. Using the gene weights of the overlapping genes in both mTCs, pair-wise correlations were calculated. Specifically, Spearman correlations were performed in R using the *pspearman*-package (v0.3-0) in R, with a t-distribution approximation to determine the P-value. As the number of genes with an absolute weight above 3 was different for every mTC, the size of the overlap in genes between two mTCs changed. The significance of the Spearman correlation found between two mTCs, therefore, was dependent on the number of overlapping genes. Hence, the significance of the found size of the overlap in genes between mTCs should be determined. To this end, for a pair of mTCs, two sets of random gene identifiers were selected from all possible gene identifiers. The amount of randomly selected genes per set corresponded to the number of genes with a weight >3 in both mTCs. Subsequently, the overlap in gene identifiers between the two random sets of gene identifiers was determined. By repeating this 10,000 times, the chance of finding a given overlap between two sets of genes could be determined.

Ultimately, mTCs were said to be concordant when their correlation was > 0.5, with a P-value < 0.05, given that there was a significant overlap in genes (P-value of overlap <0.05).

### Clustering of Metabolic Transcriptional Components, Genes and Samples

For each of the four datasets, the mixing matrix (MM) containing activity scores was clustered both on samples and mTCs. To this end, hierarchical clustering was performed using ward-D2 as the method and 1-cor(data) as the distance function. Heatmaps were created using R’s *gplots* package (v3.0.1). Based on the MM clustering for every dataset, metabolic subtypes were defined. To determine the sizes of clusters of samples that would make up a metabolic subtype, the dendrograms resulting from hierarchical clustering of the samples were systematically cut at dissimilarity values ranging from 0.0 to 8.0 with increments of 0.2. For each of the four datasets GEO, TCGA, CCLE, and GDSC, the cutoff was chosen at the dendrogram height at which the smallest cluster reached a size of 50 samples (**Figure S6**).

### CIBERSORT

Relative and absolute immune fractions for 22 immune cell types were estimated for all samples in GEO and TCGA datasets using the CIBERSORT algorithm running with default parameters, 1000 permutations, and selecting ‘absolute nosumto1’ as output. This output was then associated with the activity of the mTCs through spearman correlation.

## Statistical Analyses

Univariate OS on breast cancer samples from GEO and univariate DRFS analyses on melanoma samples from TCGA were performed using a cox regression model through *survminer* (v0.4.3) and *survival* (v2.43-3) packages in R. Confidence intervals were set at 0.95. Significance was tested through the Log Rank test. Scripts are available at github.com/**MetabolicLandscape/.** Pearson correlations were performed in R using the cor.test()-function from the *stats* package (v.3.5.1). Spearman correlations and the corresponding exact P-values were calculated using the *pspearman*-package (v0.3-0) in R, with a t-distribution as an approximation.

## ACKNOWLEDGMENTS

This research was supported by grants awarded by the Young Academy Groningen (to R.S.N.F., M.T.C.W., and V.C.L.), the Netherlands Organization for Scientific Research (NWO-VENI grant 916-16025 to R.S.N.F), the Dutch Cancer Society (RUG 2013-5960 to R.S.N.F, RUG 2014-6691 to S.J., Young Investigator Grant 10913/2017-1 to M.J.), the European Union through the Rosalind Franklin Fellowship (COFUND project 600211 to M.T.C.W.), and a grant from the Hanarth Fonds, The Netherlands (2019N1552 to R.S.N.F).

## AUTHOR CONTRIBUTIONS

R.S.N.F, V.C.L., C.G.U, and A.B collected and compiled the data. R.S.N.F., V.C.L., and C.G.U performed data analyses. All authors contributed to the data interpretation, writing of the manuscript, and the final decision to submit the manuscript.

## SUPPLEMENTARY INFORMATION

Contains supplementary Figures S1 – S8 and their legends.

To accommodate the editorial process, and due to file constraints, Supplementary Tables S1 – S5 are available as excel files but omitted from the generated composite pdf file.

---

**Figure S1 – Related to Figure 1.**
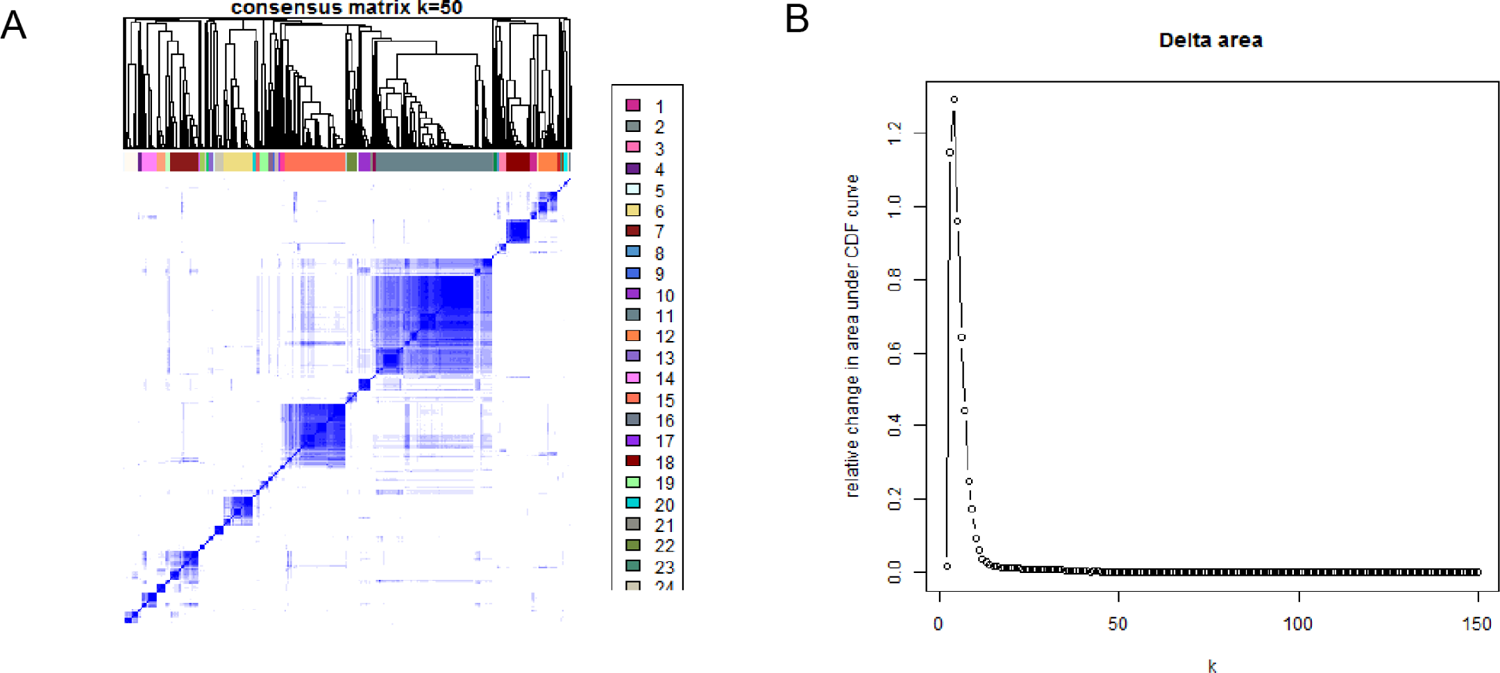
**(A)** Consensus clustering gene set enrichment scores of all TCs in the GEO dataset. Consensus matrix for a *k* of 50 gene set clusters. **(B)** Consensus clustering gene set enrichment scores of all TCs in the GEO dataset. Relative change in area under the consensus CDF curve with increasing *k*.

**Figure S2 – Related to Figure 1.**
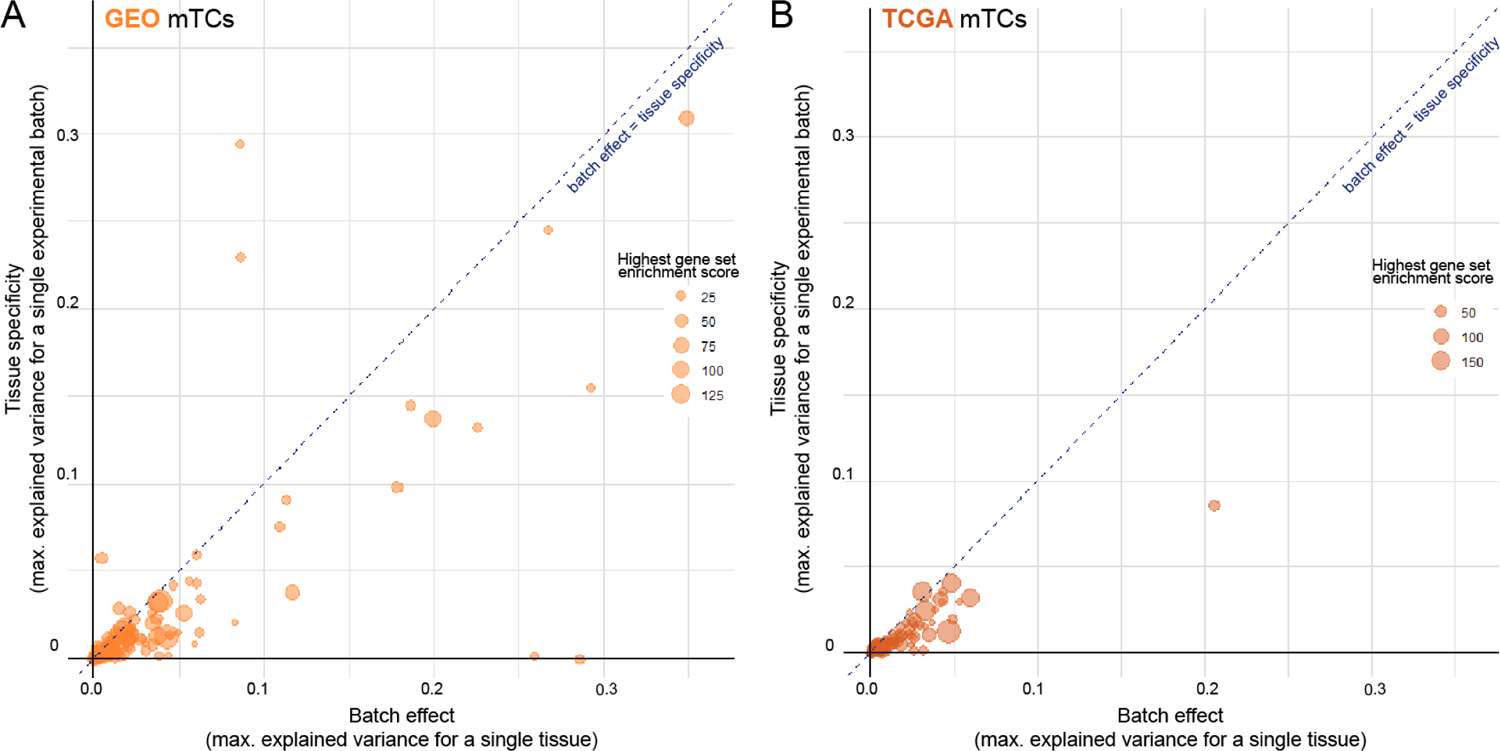
Scatter plots showing the maximum batch effect and tissue specificity for GEO **(A)** and TCGA **(B)** mTCs. Size of the dots correspond to the highest gene set enrichment score of that mTC. The magnitude of the batch effect in an mTC is estimated by the maximum fraction of the sample variance in an experimental batch that is explained by that mTC. Similarly, the tissue specificity of an mTC is estimated by the maximum fraction of the sample variance in a tissue type that is explained by that mTC.

**Figure S3 – Related to Figure 1.**
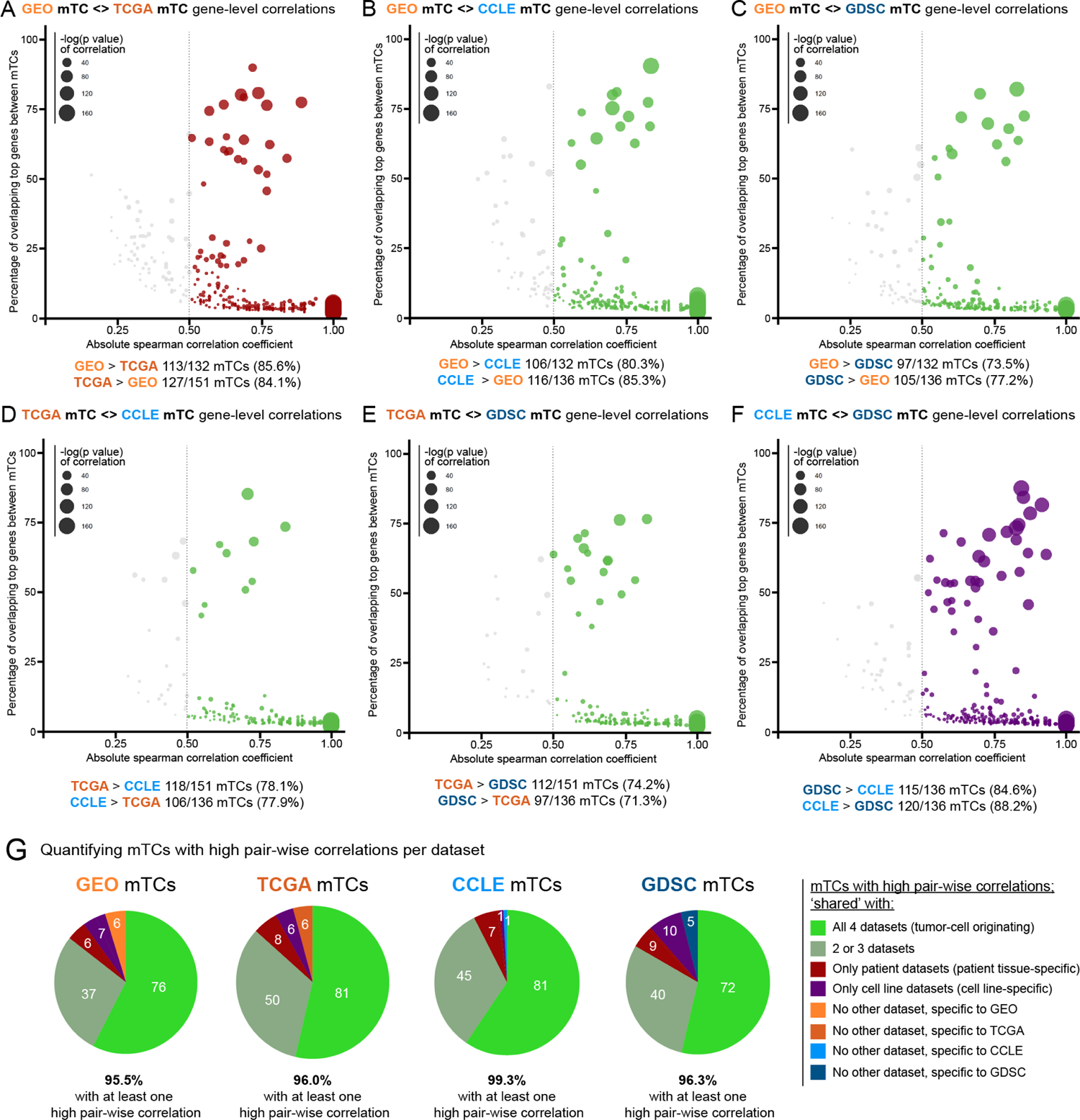
Scatter plot showing absolute spearman correlation coefficients (x-axis), versus the percentage of overlapping top genes (genes with absolute weight >3) between mTCs from different datasets (y-axis). Only significant pair-wise correlations (with P-value <0.05 and top gene overlap significance <0.05) are shown. Colored dots show absolute correlations > 0.5, the size of the dots represent the P-value of these spearman correlations. Scatter plots are shown for correlations between **(A)** GEO and TCGA mTCs, **(B)** GDSC and CCLE mTCs, **(C)** GEO and GDSC mTCs, **(D)** TCGA and GDSC mTCs, **(E)** GEO and CCLE mTCs, **(F)** TCGA and CCLE mTCs. **(G)** Pie graphs quantifying the amount of mTCs with high correlations for every dataset.

**Figure S4 – Related to Figure 2.**
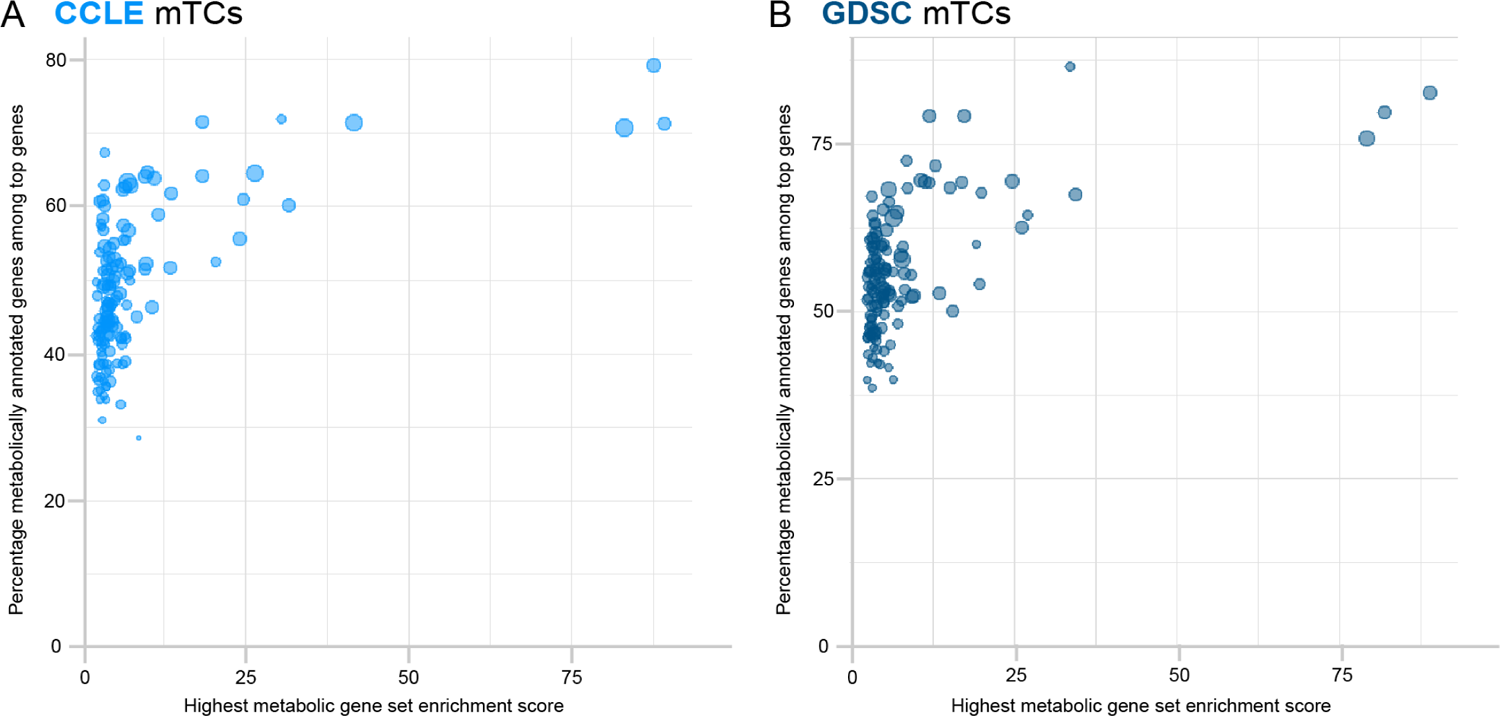
Dot plots showing the highest metabolic gene set enrichment score for every CCLE (A) and GDSC (B) mTC (x-axis) versus the percentage of metabolically annotated genes in the top genes (genes with absolute weight >3) in those mTCs (y-axis).

**Figure S5 – Related to Figure 3.**
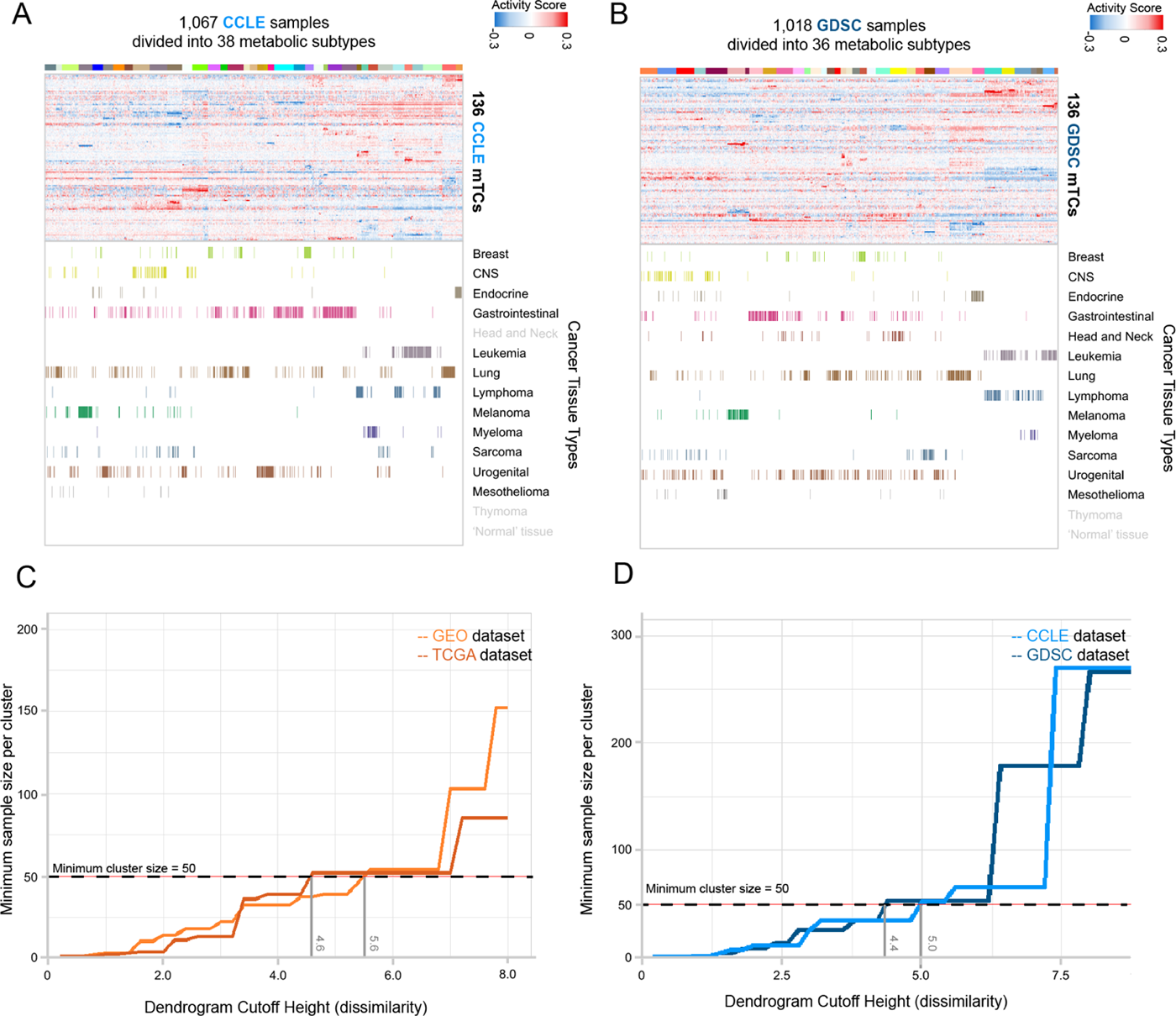
**(A)** Metabolic landscape for CCLE samples. The 1,067 samples were hierarchically clustered and divided into 38 metabolic subtypes. **(B)** Metabolic landscape for GDSC samples. The 1,018 samples were hierarchically clustered and divided into 36 clusters metabolic subtypes. Grey labels designate tissue types that are present in other datasets, but are not present in the given dataset. **(C)** Hierarchical clustering of activity scores of mTCs in samples from GEO and TCGA datasets used in order to define metabolic subtypes. The plot shows the minimum sample size of a cluster depending on the chosen cutoff height of the dendrogram resulting from hierarchical clustering. The heights at which the minimum cluster size reaches 50 is given for both GEO and TCGA datasets. **(D)** Hierarchical clustering of activity scores of mTCs in samples from CCLE and GDSC datasets used in order to define metabolic subtypes. The plot shows the minimum sample size of a cluster depending on the chosen cutoff height of the dendrogram resulting from hierarchical clustering. The heights at which the minimum cluster size reaches 50 is given for both CCLE and GDSC datasets.

**Figure S6 – Related to Figure 3.**
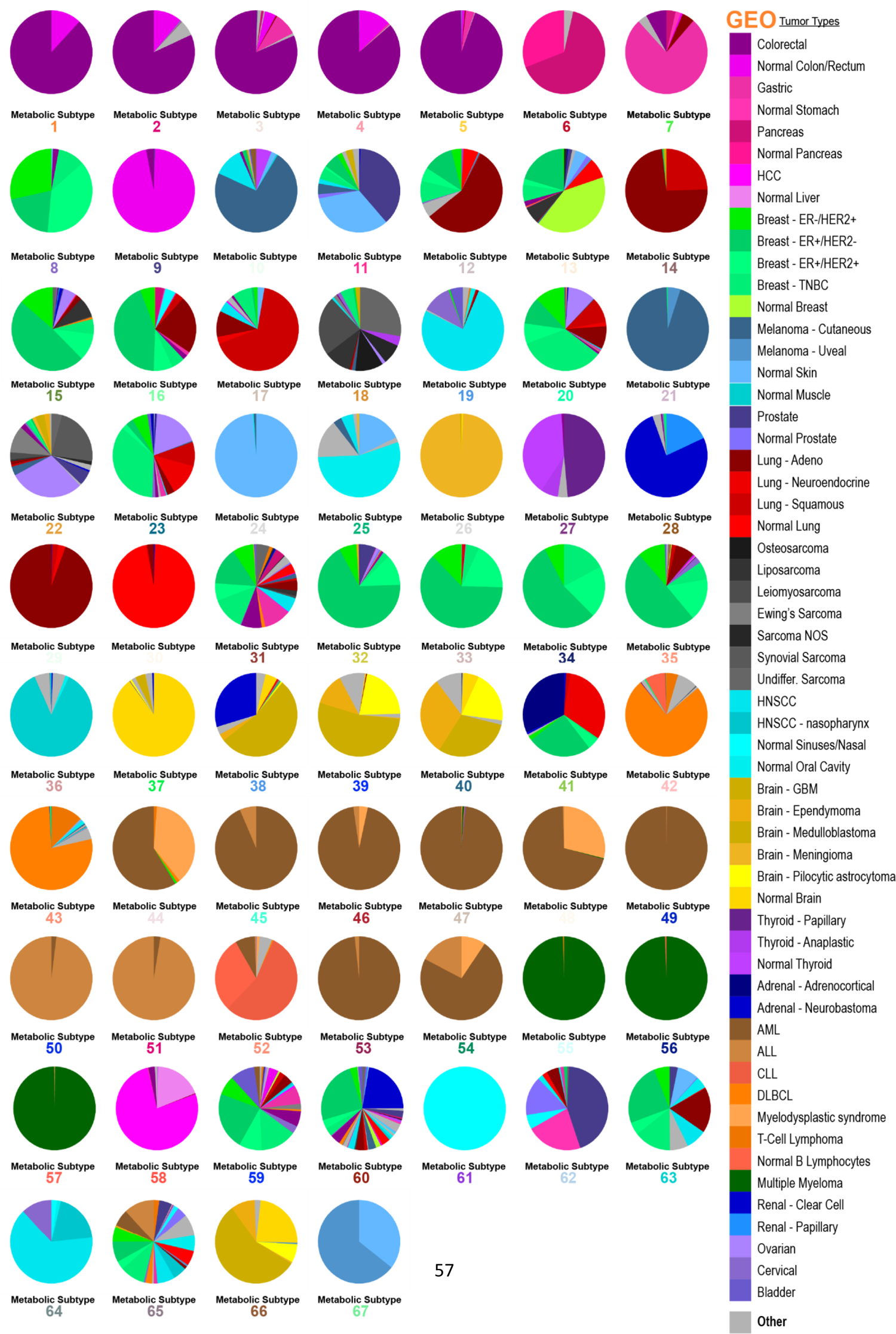
Pie graphs depicting the tissue type composition of the 67 metabolic subtypes defined for the GEO dataset.

**Figure S7 – Related to Figure 3.**
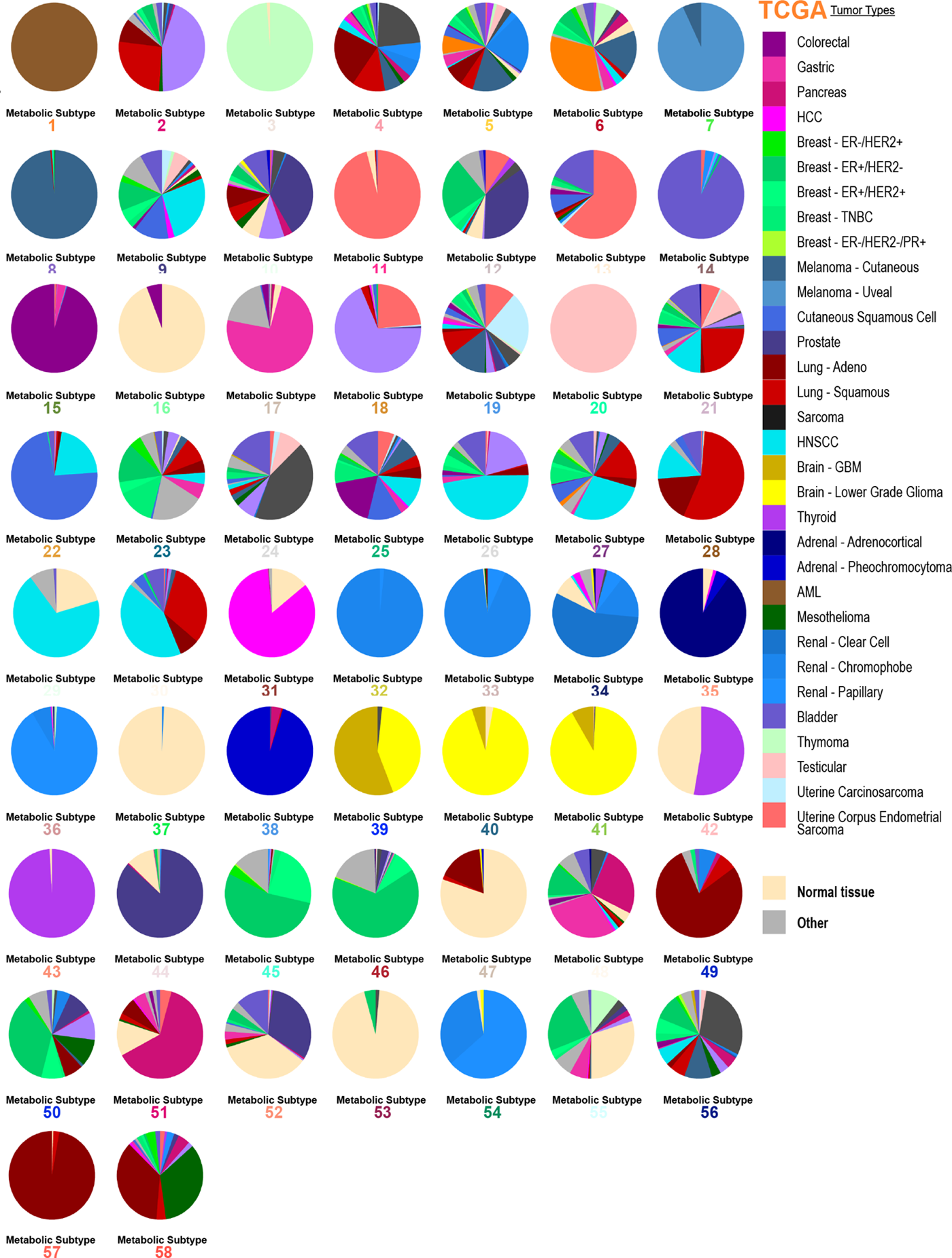
Pie graphs depicting the tissue type composition of the 58 metabolic subtypes defined for the TCGA dataset.

**Figure S8 - Related to Figure 5.**
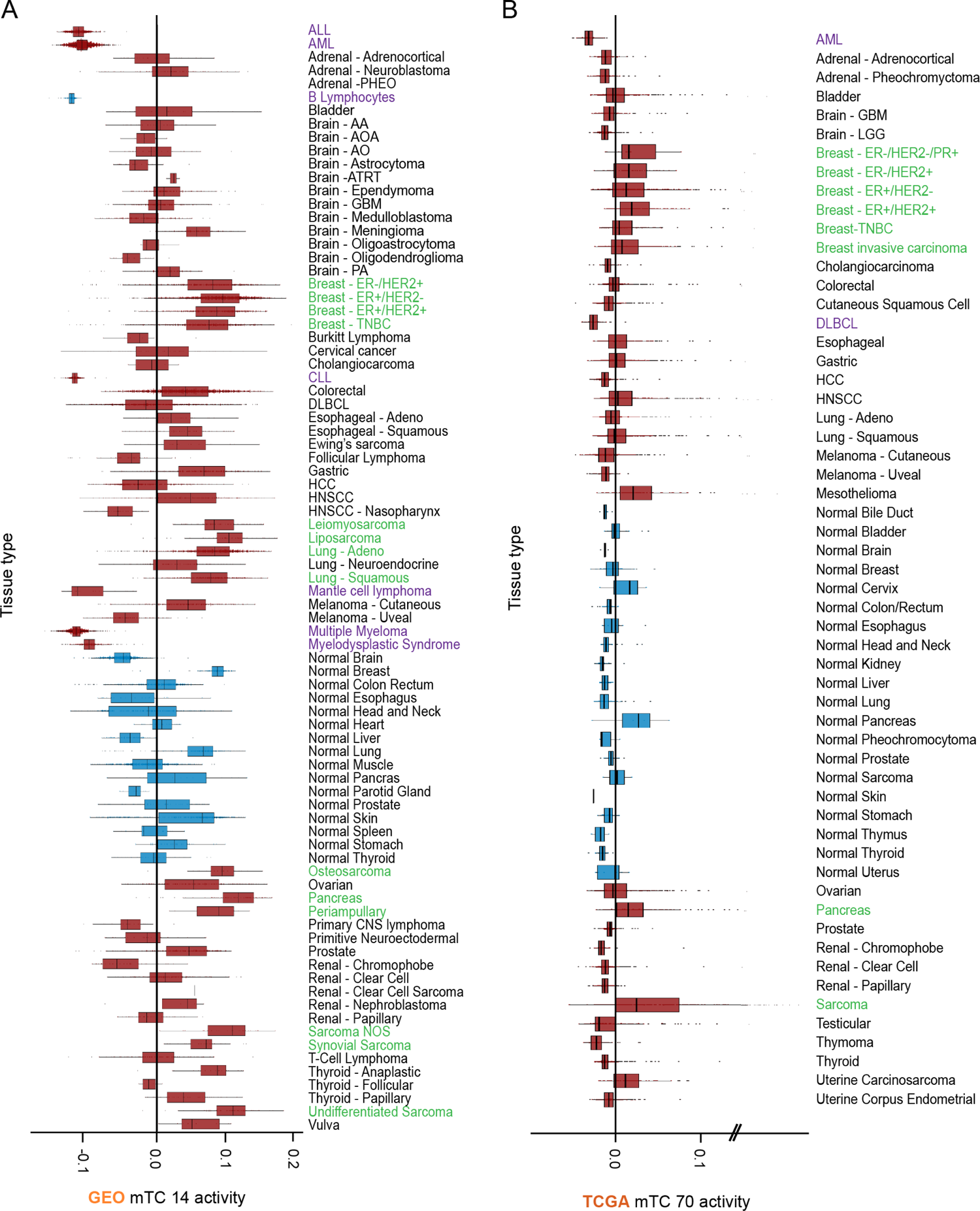
**(A)** Activity of GEO mTC 14 in samples, grouped per tissue type. Tissue types with a higher median activity highlighted in the text are given a red axis label, tissue types with a lower median activity highlighted in the text are given a blue axis label. **(B)** Activity of TCGA mTC 70 in samples, grouped per tissue type. Tissue types with a higher median activity highlighted in the text are given a red axis label, tissue types with a lower median activity highlighted in the text are given a blue axis label.

